# Repeated translocation of a gene cassette drives sex chromosome turnover in strawberries

**DOI:** 10.1101/163808

**Authors:** Jacob A Tennessen, Na Wei, Shannon Straub, Rajanikanth Govindarajulu, Aaron Liston, Tia-Lynn Ashman

**Author notes:** Current address: Department of Biology, Hobart and William Smith Colleges, Geneva, NY 14456. Current address: Department of Biology, West Virginia University, Morgantown, WV, 26506. For correspondence: Tia-Lynn Ashman, 211 Clapp Hall, 4249 Fifth Avenue, Pittsburgh, PA 15260.

## Abstract

Turnovers of sex-determining systems represent important diversifying forces across eukaryotes. Shifts in sex chromosomes, but conservation of the master sex-determining genes, characterize distantly-related animal lineages. Yet in plants, where separate sexes have evolved repeatedly and sex chromosomes are typically homomorphic, we do not know whether such translocations drive turnovers within closely related groups. This phenomenon can only be demonstrated by identifying sex-associated nucleotide sequences, still largely unknown in plants. The wild North American octoploid strawberries (*Fragaria*) exhibit separate sexes (dioecy) with homomorphic, female heterogametic (ZW) inheritance, yet sex maps to at least three different chromosomes. To characterize these turnovers, we sequenced the complete genomes of 60 plants of known sex from five *Fragaria* taxa. We identified 31-mers unique to females and assembled their reads into contigs. Remarkably, a short (13 kb) sequence is observed in nearly all females and never in male-fertile individuals, implicating it as the sex-determining region (SDR). This female-specific “SDR cassette” contains both a gene with a known role in fruit and pollen production and a novel retrogene absent on Z and autosomal chromosomes. Comparing SDR cassettes across taxa reveals a history of repeated translocation, which can be ordered temporally due to the capture of adjacent sequence with each successive move. The accumulation of these “souvenirs” suggests an adaptive basis for the expanding (up to at least 23 kb) hemizygous region. This is the first plant SDR known to be translocated, and it suggests a new mechanism (“move-lock-grow”) for expansion and diversification of incipient sex chromosomes.

**Significance Statement:** Sex chromosomes frequently restructure themselves during organismal evolution, often becoming highly differentiated. This dynamic process is poorly understood for most taxa, especially during the early stages typical of many dioecious plants. In wild strawberries, a sex-determining region of DNA has repeatedly changed its genomic location, each time increasing the size of the hemizygous female-specific sequence. This observation shows for the first time that plant sex regions can “jump”, and suggests that this phenomenon may be adaptive by gathering and locking new genes into linkage with sex. This conserved and presumed causal sequence with a variable genomic location presents a unique opportunity to understand how sex chromosomes first begin to differentiate.

## Introduction

Sex chromosomes are one of the most dynamic components of eukaryotic genomes [1]. The defining feature of a sex chromosome, the sex-determining region (SDR), has experienced similar restructuring in multiple independent instances of autosomes evolving into heteromorphic sex chromosomes [2]. Specifically, loci under sexually antagonistic selection become linked to the SDR, recombination is suppressed, and the hemizygous region grows in size [3]. The mechanisms of this chromosome restructuring may involve successive inversions of the SDR or translocations of sequence on and off the sex chromosome [3,4]. Turnovers that change the genomic location of the SDR have been revealed in the evolution of animal sex-determining systems [1,5-8], where they may be important drivers of sexual dimorphism and speciation [9,10]. While theory on the processes driving these transitions is growing [11-14], few systems exist in which the mechanisms of turnovers can be inferred [15-17].

Fundamental questions about SDR turnovers thus remain unanswered. Do turnovers typically involve mutations in new loci that take control of an existing sex-determining mechanism [18,19], functionally independent mutations [20], or translocations of the existing sex-determining gene(s) [21-24]? Similarly, do turnovers typically restart the process of SDR divergence, maintaining “ever young” sex chromosomes [25], or do they contribute to increasing chromosome heteromorphy via loss or gain of sequence [11,14,26]? And ultimately, is there an adaptive basis for these turnovers? Although master sex-determining genes like *SRY* and *DMRT1* are highly conserved in some animal systems, the causal SDR loci or gene cassettes remain unknown for most dioecious eukaryotes [27]. Even less is known about the temporal order of turnovers in any taxon, and thus directional trends in sex chromosomal rearrangement [2].

Turnovers of SDRs are likely to be quite common in plants, given that dioecy (separate males and females) has evolved repeatedly from hermaphroditism (combined male and female function) and most sex chromosomes are young and homomorphic [28,29]. Yet despite the potential of dioecious plants for yielding evolutionary insights, there are disproportionately few systems with mapped SDRs [28,29], or known causal genes [30], and the pattern or mechanism of turnovers remain entirely unexplored. The octoploid strawberries (*Fragaria*; 2*N* = 56; four disomically-inherited subgenomes AvBiB1B2) show recently evolved dioecy with substantial genetic and phenotypic diversity in sexual system among closely related taxa, and thus have been a model system for studying incipient plant sex chromosomes [31-35]. These species all possess homomorphic ZW chromosomes with male sterility strongly correlated with female fertility [34], and with a single SDR explaining the majority of variation in male and female function, though the degree of sexual dimorphism varies across taxa [32,36-39]. The SDR has been mapped to three distinct subgenomes in three geographically distinct taxa: (from eastern to western North America) *F. virginiana* ssp. *virginiana* [36], *F. virginiana* ssp. *platypetala* [35], and *F. chiloensis* [34,37] (Fig. 1). These three SDRs occur in unique sections of different chromosomes from the same homoeologous group (Fvb6_0-5.5 on B2, Fvb6_13 on B1, and Fvb6_37 on Av; notation throughout: chromosome on diploid *F. vesca* reference genome *Fvb* followed by position in Mb [Fvb1-7_Mb] [40]). The autosome Fvb6 may possess sexually antagonistic genes that predispose it to become a sex chromosome [41], as seen in other systems [42-44]. Genetic maps of *F*. ×*ananassa* ssp. *cuneifolia*, a natural hybrid of two of the taxa (*F. virginiana* ssp. *platypetala* and *F. chiloensis*), corroborate the two SDR locations of the two progenitor species [45]; other subspecies including *F. virginiana* ssp. *glauca* have yet to be studied in detail. Thus, *Fragaria* provides a unique opportunity to test whether sex chromosome turnovers represent translocations of the same SDR.

**Figure 1.**
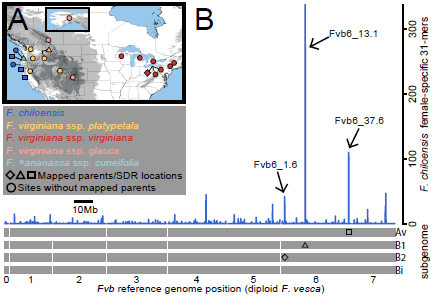
Collection localities, SDR map locations, and female-specific 31-mers aligned to *Fvb* reference genome. (A) Five North American taxa were collected (Table 1). (B) Female-specific 31-mers in *F. chiloensis* were clustered in several narrow (100 kb) genomic windows, including the three known male-sterility regions on Fvb6 (arrows). Male sterility in *F. chiloensis* has been repeatedly mapped between Fvb6_37.428-37.708 Mb on subgenome Av (square) [34,37]. Malesterility in *F. virginiana* ssp. *platypetala* and *F*. ×*ananassa* ssp. *cuneifolia* had been mapped between Fvb6_12.935-13.355 Mb on subgenome B1 (triangle) [35,45]. Male sterility in *F. virginiana* ssp. *virginiana* was previously mapped to subgenome B2 [36,42] and here was fine-mapped between Fvb6_ 1.630-1.770 Mb (diamond).

Here we use whole genome sequencing and molecular evolutionary analysis to characterize SDRs on different chromosomes across multiple octoploid *Fragaria* taxa. Our goal is to determine whether or not sex chromosome diversity reflects a single W chromosome-specific sequence that has translocated among genomic locations. Remarkably, we find an “SDR cassette” shared by females across taxa and never in male-fertile plants, containing two putatively functional genes. The SDR cassette has translocated at least twice, bringing along adjacent sequence that contributes to a widening hemizygous region and reveals the translocation order and evolutionary history of the SDRs. Thus, we report the first case of a repeatedly translocating SDR in plants, and propose a new hypothesis for sex chromosome differentiation.

## Results and Discussion

### Conserved, translocated female-specific sequence across taxa

Demonstration of SDR translocation begins with identifying homologous sex-linked sequence across taxa [27]. We identified sequence unique to the W chromosome by sequencing the complete genomes of 31 females and 29 male-fertile plants in five taxa (Table 1 and Fig. S1; range of coverage relative to the haploid reference genome = 16–57×; median = 33×). These represented the North American range of the octoploid *Fragaria* and included the parents of several previous mapping crosses for male sterility [31,34,35,37,45] (Fig. 1*A*). In the three taxa for which we had at least 9 female and 8 male-fertile plants, *F. virginiana* ssp. *virginiana,F. virginiana* ssp. *platypetala*, and *F. chiloensis*, we observed 1215, 468, and 1528 female-specific 31-mers, respectively. We identified these 31-mers with an alignment-free approach [46] but most could be subsequently aligned to *Fvb* (Figs. 1*B* and S2). A particularly marked pattern was observed for *F. chiloensis*: 38% of all female-specific 31-mers occurred in five 100 kb windows on *Fvb*, each with at least 42 female-specific 31-mers aligning at: Fvb4_21.3, Fvb6_1.6, Fvb6_13.1, Fvb6_37.6, and Fvb7_18.5 (Fig. 1*B*). Notably, three of these windows correspond to mapped male-sterility positions across taxa (Fig. 1*B*), suggesting translocation of sequence shared by the current *F. chiloensis* SDR and these *Fvb* windows. All 27 females of these three taxa across their North American range possessed 31-mers overlapping the same site homologous to Fvb6_1.636, but with a 23 bp “diagnostic deletion” not seen in the diploid hermaphrodite *F. vesca* or any male-fertile plants. The diagnostic deletion also occurs in the single *F.* ×*ananassa* ssp. *cuneifolia* female and one of the three *F. virginiana* ssp. *glauca* females. The two (out of 31) females lacking the diagnostic deletion could possess nonhomologous SDR(s) or simply carry distinct versions of this sequence. Thus, 94% of all the females across the five taxa have a shared SDR association with the Fvb6_1.6 region, from which sequence has repeatedly translocated to the multiple mapped locations.

**Table 1.**
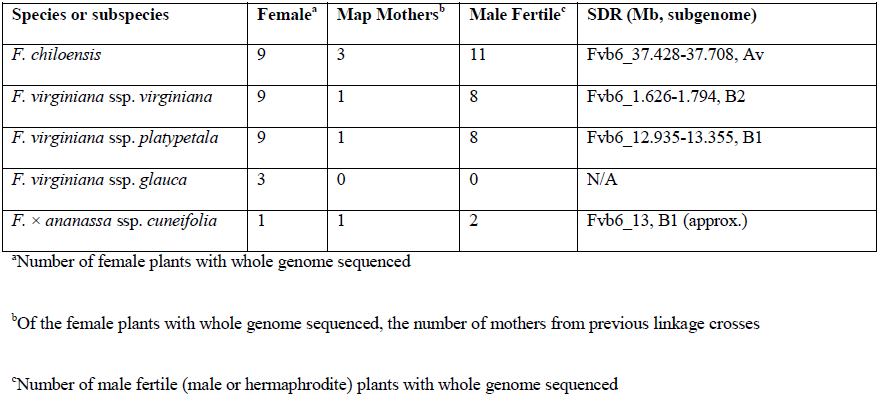
*Fragaria* taxa sequenced and SDR positions mapped in linkage crosses

Reconstructing the history of W-specific sequence requires precisely established SDR map locations across taxa, as well as concrete knowledge of the orthologous Z and autosomal sequences that serve as a proxy for the ancestral state. The SDR locations at Fvb6_13 on B1 [35] and Fvb6_37 on Av [34,37] were previously mapped to within 300 kb (Fig. 1*B*). However, the SDR location in *F. virginiana* ssp. *virginiana* was only known within 5.5 Mb from an F1 cross [36,41]. We thus fine mapped male sterility in this same cross (*N* = 1878) to a 168 kb region between Fvb6_1.6-1.8 on B2. Next, we assembled maternal-parent bacterial artificial chromosomes (BACs) overlapping this region, and used Fluidigm genotyping to further fine map male sterility between Fvb6_1.630 and Fvb6_1.770 (Fig. 1*B*). Variants in repulsion with male sterility allowed us to assign two BAC scaffolds to the Z chromosome, but no BACs overlapping this revised male sterility region were recovered from the W chromosome itself (Fig. S3). This fine-mapped region matched a spike in female-specific 31-mers seen in other taxa (Figs. 1*B*, S2), indicating homology to a widespread sex-associated sequence.

### Stratified souvenir sequences reveal translocation history

Patterns of shared and distinct sequences at the SDR, combined with phylogenetic and linkage mapping data, allow us to reconstruct the details of its history to an unprecedented degree. We assembled a 2.7 kb haplotype (0-30% missing data; mean = 16%) overlapping the diagnostic deletion for all females (*N* = 29) except the two *F. virginiana* ssp. *glauca* females lacking it. The phylogeny of these sequences resolved three distinct clades, α, β, and γ (Fig. 2). Remarkably, each clade was associated with a distinct SDR map location, thus revealing their evolutionary progression. The α clade contained females of *F*. *virginiana* ssp. *virginiana, F*. *virginiana* ssp. *platypetala*, and *F*. *virginiana* ssp. *glauca,* including the mother for which male sterility maps to Fvb6_1 (see above). The β clade contained females of *F*. *virginiana* ssp. *platypetala* and *F.* ×*ananassa* ssp. *cuneifolia*, including two for which male sterility maps to Fvb6_13 [35,45]. The γ clade contained females of *F*. *virginiana* ssp. *virginiana* and *F*. *virginiana* ssp. *platypetala*, as well as a monophyletic group comprising all nine *F. chiloensis* females including the three for which male sterility maps to Fvb6_37 [34,37]. Most *F*. *virginiana* ssp. *virginiana* plants were in the α clade, while *F. virginiana* ssp. *platypetala* was nearly even distributed across clades. Because the β and γ clades are sister to each other with strong support (93% SH-like; 92% bootstrap; Fig. 2), at least one of their map locations (Fvb6_13 and/or Fvb6_37) must be newly derived relative to the α clade, consistent with an ancestral map location at Fvb6_1.6, the only region homologous to sequence shared across all three clades.

**Figure 2.**
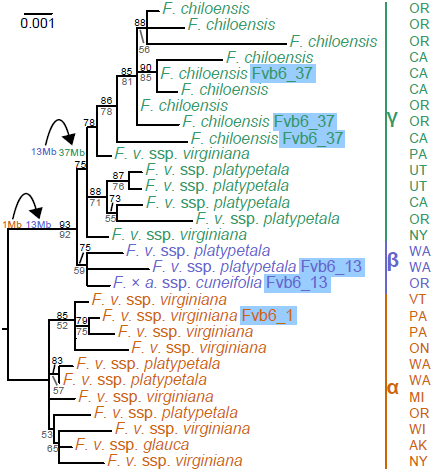
Phylogeny of central 2.7 kb of SDR cassette, including the diagnostic deletion and *RPP0W*. All females, with species or subspecies indicated, occur in one of three major clades, α, β, and γ. U.S. state or Canadian province of origin is shown on right. Support for clades, if greater than 50%, is indicated as SH-like support (above branches, black) or bootstrap support (below branches, grey). For maternal parents in linkage crosses, the map position of their SDR is highlighted with a blue box. Each of the three major clades is associated with a different map location: Fvb6_1, Fvb6_13, or Fvb6_37. Outgroups (not shown) are autosomal paralogs. The close evolutionary relationship between the β and γ clades is consistent with the inferred history of translocations, indicated at two points with black curved arrows to the left of the phylogeny.

The *F. chiloensis* monophyly allowed us to assemble a 28 kb haplotype containing 89% of the female-specific 31-mers for this species (Fig. 3, top colored bar). This assembly was guided by the 5109 31-mers present in at least eight *F. chiloensis* females and still absent in all male-fertile plants. We first assembled female-specific sequence homologous with the Fvb6_1.6 region, generating three contigs comprising 13 kb. These contigs were ordered and oriented into a unified haplotype, the SDR cassette, based on homology with Z-chromosome BACs from *F. virginiana* ssp. *virginiana* (for *F. chiloensis*, 98% sequence similarity across 10.4 kb aligned), and with diploid *F. vesca* at Fvb6_1.635-1.642 (for *F. chiloensis*, 93% sequence similarity over 8.6 kb aligned) (Fig. 3). This SDR cassette was nested within an additional 10 kb of sequence on either side (the “flanking” sections) that included 5 kb homologous to Fvb6_13.1 consistent with the β clade SDR map location [35], as well as 2 kb homologous to Fvb4_21.3. These sections were nested within an additional 5 kb of sequence primarily showing homology to Fvb6_37.6 (the “outer” section), consistent with the γ clade *F. chiloensis* SDR map location [34]. An additional 7% of the *F. chiloensis* female-specific 31-mers do not align to this haplotype but are presumably closely adjacent, aligning to Fvb6 between 37.59-37.61. Thus, the *F. chiloensis* SDR at Fvb6_37.6 encloses nested “souvenir” sequence matching the known SDR locations in other taxa, consistent with a history of movement from those locations.

**Figure 3.**
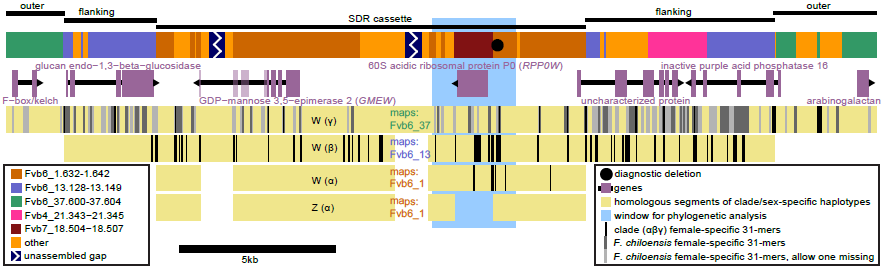
Reconstructed W-specific haplotypes. Starting at the top, the “SDR cassette,” “flanking”, and “outer” sections of the 28 kb haplotype reconstructed from *F. chiloensis* are shown, followed by the haplotype colored according to homology with reference genome *Fvb*. The two assembly gaps and the diagnostic deletion are indicated on this color-coded haplotype. Predicted genes are shown just below with direction of transcription indicated by arrowheads; *GMEW* exons after the nonsense polymorphism are faded. Beneath those are the inferred homologous portions of the W-specific haplotypes from each clade highlighting the portions containing female-specific 31-mers. They are followed by the Z-specific sequence obtained from BACs of a maternal *F. virginiana* ssp. *virginiana* linkage cross parent. The light blue rectangle indicates the 2.7 kb window used in phylogenetic analysis (Fig. 2). All three clades share the SDR cassette, suggesting it is the oldest and that Fvb6_1.6 is the original SDR position. Clades β and γ also share the flanking sections, suggesting a translocation to Fvb6_13.1. Only clade γ possesses the outer section, consistent with a second translocation to Fvb6_37.6 unique to this clade.

To assess the evolutionary history of the SDR we characterized in detail the sequence neighboring the SDR cassette in each of the three clades (Figs. 2 and 3). Female-specific 31-mers within each clade (but absent in all 29 male-fertile plants) aligned to distinct portions of the assembled *F*. *chiloensis* W haplotype (Fig. 3, vertical black lines). Interestingly the 31-mers from the α clade were restricted to the SDR cassette, specifically all within 1.7 kb of the diagnostic deletion, consistent with the SDR remaining at the ancestral Fvb6_1.6 location within this clade [36,41]. The 31-mers from the β clade, on the other hand, occurred in both the SDR cassette and flanking sections, suggesting the SDR had jumped from Fvb6_1.6 to Fvb6_13.1 in the ancestor of the β and γ clades (Figs. 2 and 3). A second translocation to Fvb6_37.6, specific to the γ clade, explains the SDR cassette and flanking sections retained in γ from its previous locations and also the outer sections unique to γ with homology to Fvb6_37.6, as well as the map location at Fvb6_37.6 in three γ mothers (Figs. 2 and 3) [34,37]. Thus, as translocations carried “souvenir” sequence from each of their previous locations, we can for the first time ascertain the temporal order of SDR movements (Fig. 2). It is possible that there have been additional translocations to not-yet-observed SDR locations, perhaps still segregating in these or other populations. In addition, a 2 kb portion of the downstream flanking section shows homology to Fvb4_21.3, which could be a souvenir from another prior SDR location, or an independent translocation of sequence into the SDR, commonly seen in sex chromosomes [2]. Intriguingly, in the diploid *F. vesca* ssp. *bracteata*, the gynodioecious cytoplasm donor to the *Fragaria* octoploids, dominant male sterility maps to Fvb4 [47], though 9 Mb away at Fvb4_30. The repeated translocations occurred relatively rapidly: octoploid *Fragaria* only originated approximately 1 mya [48], SDR clades share high (> 99%) sequence similarity (Fig. 2), and all three SDR clades still co-exist within *F. virginiana* ssp. *platypetala*, which occurs between the other taxa geographically (Fig. 1*A*) and phylogenetically [49]. This unparalleled insight into the temporally-ordered series of SDR translocations in octoploid *Fragaria* is the first of its kind in any plant or animal system, and as such will provide a unique opportunity to examine fitness effects of SDR location.

Many flowering plant SDRs are large or coarsely mapped, complicating the identification of causal genes [28,29]. In contrast, only two coding genes, annotated as GDP-mannose 3,5-epimerase 2 (here *GMEW*) and 60S acidic ribosomal protein P0 (here *RPP0W*), were identified in the SDR cassette (Fig. 3). *GMEW* homologs occur on the Z chromosome BAC (99% similarity) and at Fvb6_1.6 (98% similarity). In clades β and γ but not α, *GMEW* has a premature stop codon shortening the coding sequence from 376 to 222 residues. GDP-mannose 3,5-epimerase converts GDP-mannose to GDP-L-galactose in vitamin C and cell wall biosynthesis [50,51], affecting fruit development in *Fragaria* [52,53] and pollen production in other plants [51]. While *GMEW* is a plausible candidate, the polymorphic stop codon may suggest a variable role among females. *RPP0W* falls within a 1.2 kb W-specific insertion and shows 99% similarity to a gene at Fvb7_18.5, but lacks that gene’s four introns, suggesting it is retrotransposed. Ribosomal proteins are essential for polypeptide synthesis and are often retrotransposed [54]. In plants they can affect processes from development to stress response [55], with mutations sometimes acting dominantly [56], as expected for the first mutation in a female heterogamic (ZW) system [1]. In rice, the overaccumulation of ubiquitin fusion ribosomal protein L40 results in defective pollen and male sterility [57]. In classic two-gene SDR models, one gene affects male function and another female function [58], though a single master regulator could also perform both roles [30]. The diagnostic deletion and the repetitive unassembled gaps, though apparently noncoding, could also be functional motifs. Our BLAST annotation did not identify any transposons, and the only plant repetitive sequences identified within the assembled haplotype were stretches of dinucleotide repeats under 50 bp. In *F. vesca*, both *RPP0W* and *GMEW* homologs show decreasing expression during anther development and even lower expression within pollen [59], but expression profiles in octoploids remain to be characterized. Neither gene family (of *GMEW* nor *RPP0W*) has been directly implicated in sex determination, highlighting that many pathways could alter plant sex function [60,61].

### Increasing size of SDR births a new adaptive hypothesis

Turnovers of SDR locations are common in evolution [4-6,10,11], suggesting an adaptive basis for these rearrangements. Translocations could be favored to escape genetic load [13] or to acquire linkage with loci under sexually antagonistic selection [14]. The *Fragaria* SDR cassette translocations are consistent with either adaptive scenario, as fitness effects of the genomic neighborhoods remain unknown. In addition, a third adaptive explanation is suggested by the observation that each jump of the SDR increased the size of the hemizygous female-specific haplotype by accumulating souvenir sequences. The α clade is only hemizygous for the 1.2 kb insertion containing *RPP0W*. The β clade is hemizygous for the 13 kb SDR cassette and its two genes are maintained in perfect linkage disequilibrium with sex. The γ clade is hemizygous for the 23 kb of SDR cassette and flanking sections containing five genes in perfect linkage disequilibrium with sex. Adjacent non-coding sequence could also be functional. If SDR genes are under sexually antagonistic selection, as seen for some *Fragaria virginiana* traits [38], then a benefit of translocation is to “lock” them into hemizygosity, and translocation *per se* is adaptive. Thus, these results could represent a previously unrecognized mechanism of sex chromosome evolution – “move-lock-grow” – which could explain the rapid differentiation and dynamic genomic rearrangements of incipient SDRs. Intriguingly, *F. chiloensis* shows greater male-female differences than other *Fragaria* [37] as well as sex-specific recombination rates [34], and is fixed for γ in our samples (Fig. 2), whereas *F. virginiana* shows less pronounced and more variable sexual differentiation [38,39] and harbors all three clades with α the most common (Fig. 2). This is consistent with a correlation between SDR size/content and sexual dimorphism. A similar growth mechanism may underlie other hemizygous supergenes [62].

## Conclusions

A conserved SDR cassette has repeatedly changed genomic location across octoploid *Fragaria*, supporting a translocation model of sex chromosome turnover. This is the first unambiguous evidence of SDR translocation in flowering plants, as it is rarely possible to distinguish translocations from de novo innovations unless putative causal sequences have been identified in more than one taxon [29,63]. In Salicaceae, SDRs occur on different chromosomes with no evidence of large-scale rearrangements, but data thus far are consistent with either master/slave regulatory dynamics [18] or SDR jumps [64,65]. Turnovers involving reversal of heterogamety, as seen in *Silene* [66], are more likely to be fusions of sex chromosomes to autosomes rather than translocations of SDR sequence to new chromosomes. Our discovery of a conserved yet mobile W-specific locus helps to unify extensive disparate research on the genetic basis of dioecy in *Fragaria* and across flowering plants [37,63]. It suggests that independent mechanisms of dioecy within closely related taxa may be rarer than they appear. Instead, SDR translocation can maintain the same genetic basis for sex while adjusting genomic location and accumulating sequence that may contain sexually antagonistic alleles as well as increasing recombination suppression within the growing hemizygous SDR. The “move-lock-grow” mechanism may allow for rapid and extensive change in sex chromosomes, likely generating pronounced evolutionary and ecological consequences.

## Materials and Methods

### Samples

F1 offspring from the previously described *F. virginiana* ssp. *virginiana* cross “Y33b2×O477” [36,41] were sexed (*N* = 1878) as described below and genotyped (*N* = 184) at sex-linked microsatellite markers [36] to identify possible recombinants, which were sequenced with targeted capture (*N* = 67) as previously described [40]. For the whole genome analysis, we examined 60 outbred, unrelated plants distributed across the geographic ranges of the octoploid *Fragaria* species (Table S1). These samples were collected from the wild as clones or obtained from the USDA National Clonal Germplasm Repository.

### Sex phenotyping

We determined sex using our established method [41]. In brief, we grew plants with 513 mg granular Nutricote 13:13:13 N:P:K fertilizer (Chisso-Asahi Fertilizer) under 15°:20° C night:day temperatures and 10 to 12 h days, and then exposed them to 8°:12° C night:day temperatures with an 8 h low light day to initiate flowering. Fertilizer and pest control measures were applied as needed. Male function was scored as a binary trait: plants with large, bright-yellow anthers that visibly released pollen were “male-fertile,” and plants with vestigial white or small, pale-yellow anthers that neither dehisced nor showed mature pollen were “female”. Because of the tight correlation between male function and female function, male sterility serves as a good phenotypic marker of the SDR [34].

### BAC sequencing

A BAC library was prepared by Chris Saski, Clemson University Genomics Institute (CUGI) from 90 g leaf tissue collected at the University of Pittsburgh (UPitt) from Y33b2, the female parent of the *F. virginiana* ssp. *virginiana* linkage mapping cross [36,41]. BAC construction methods followed [67] with minor modifications. We designed overgo probes from the mapped male sterility region between Fvb6_1.626-1.794 (Table S2). We labeled probes individuallywith 32_P following the CUGI protocol (http://www.genome.clemson.edu/resources/protocols) and hybridized them to the BAC filters at 60°C overnight. This yielded 69 positive clones (Table S2).

Genomic libraries from these 69 BACs were individually prepared and barcode indexed with the Illumina TruSeq DNA HT kit and sequenced with 150 bp paired end reads on a single lane of Illumina MiSeq at Oregon State University Center for Genome Research and Biocomputing (OSU CGRB). Reads were quality trimmed for both Q> 20 and Q> 30 with Trimmomatic [68] and merged, when possible, with the program FLASH [69]. We filtered merged reads and unmerged pairs by digital normalization at coverage of 100 using khmer [70]. For each library, both quality trimming sets were de novo assembled with Velvet [71] using a range of kmers from 31-91 bp. We selected the assembly with the longest contig for downstream analyses of each BAC (Table S2).

We masked vectors with bedtools [72] and used BLAT to identify identical overlap of > 1 kb among BACs. Groups of BACs representing putative homoeologs were imported into Geneious R7 [73] and further scaffolded manually. The resulting 11 assemblies were assigned to homoeologs (Fig. S3) by the presence of linkage mapped SNPs observed in the target capture and microfluidic markers. We aligned a shared 19.819 kb region with MAFFT [74] and estimated a maximum likelihood tree with PhyML [75], confirming the identification of four pairs of homologous chromosomes.

### DNA extraction and quantification

For Fluidigm genotyping, genomic DNA was extracted from silica dried leaf tissues using Norgen Biotek Plant/Fungi DNA Isolation 96-Well Kit (Ontario, Canada), and by the service provider Ag-Biotech (Monterey, CA, USA). An additional 100 μ l 10% SDS and 10 μl β - mercaptoethanol were added to the lysis buffer to improve DNA yield. DNA was further purified with sodium acetate and ethanol precipitation. DNA concentration was quantified by Quant-iT PicoGreen (Invitrogen, Carlsbad, CA, USA) assays at UPitt Genomics Research Core (GPCL).

### Amplicon fine mapping

We designed Fluidigm microfluidic markers for fine mapping Y33b2×O477 following our previous methodology [34]. We designed primer pairs for 48 amplicons with mean expected size of 385 bp: 12 and 16 on the two BAC contigs corresponding to the Z homoeolog (Fig. S3), and 20 between Fvb6_0.716-17.605 (Table S3). We used the Fluidigm 48.48 Access Array IFC (Integrated Fluidic Circuits) at University of Idaho IBEST for amplicon library preparation following standard simplex reaction protocol. We pooled the amplicons of 190 F1 offspring and the two parents for paired end 300 bp sequencing on a 1/4 lane of Illumina MiSeq. We trimmed reads as above, aligned them to *Fvb* and the BAC sequences using BWA v. 0.7.12 [76], and called genotypes with POLiMAPS [40]. We identified recombinants and used these to define the narrowest possible window overlapping male function.

### Whole genome sequencing

Genomic DNA extraction and library preparation were performed by the OSU CGRB and at UPitt. We sheared DNA to 300 bp using a Bioruptor Pico (Diagenode, Denville, NJ), and used the NEBNext Ultra DNA Library Prep Kit for Illumina (New England BioLabs, Ipswich, MA) with individually indexed dual barcodes. We sequenced whole genomes of 60 *Fragaria* samples using four lanes of paired end 150 bp on an Illumina HiSeq 3000, with 13 to 20 samples per lane (Table 1).

### Analysis of whole genome data

We converted FASTQ files to FASTA and used Jellyfish 1.0.2 [77] to count 31-mers in each sample, the largest k-mer size allowed by Jellyfish (Fig. S1). We used the Linux *sort* and *join* functions to combine lists of 31-mers and generate lists of 31-mers shared by sets of females and absent in sets of male-fertile plants. To aid the assembly of the W-specific haplotype in *F. chiloensis*, we also generated lists of 31-mers shared in all but one female, assuming that a W-specific 31-mer could be absent due to insufficient coverage or a rare sequence variant. We aligned these 31-mers to *Fvb* using BLAT v. 32×1 [78] and retained hits with at least 29bp matching and gaps no larger than 20bp. We extracted reads containing female-specific 31-mers, and their mate pairs, from the original FASTQ files. We assembled these manually in BioEdit v.7.2.5 [79], when possible guiding the assembly with alignment to homologous *Fvb* or BAC sequences. Gaps between contigs containing female-specific 31-mers were manually joined with additional reads as possible. We assembled the central 2.7 kb of the SDR cassette, including the diagnostic deletion and *RPP0W*, for all females possessing it. We assembled a pseudo-outgroup sequence based on homologous portions of *Fvb* and BAC6 and used RAxML [80] with -m GTRCAT to generate a phylogeny of the W sequence. Major clades (α, β, and γ) were assigned visually.

We assigned portions of the W haplotype to *Fvb* regions using BLAST at GDR [81] (Table S4). We identified genes using GENSCAN [82] and annotated them with BLAST to the NCBI database and to *Fvb* which is annotated, using GDR [81]. Adjacent genes (Table S5) were identified from the *Fvb* annotation. Gene expression data in *F. vesca* [54] were extracted from http://mb3.towson.edu/efp/cgi-bin/efpWeb.cgi. We looked for significant (e-value < 0.05) hits to repetitive sequence by BLASTing to the TIGR Plant Repeat Databases [83] with GDR [81].

### Data Availability

Reads from whole genome sequencing (Bioproject Accession XXXXXXXXXXX) have been uploaded to NCBI SRA. The reconstructed W haplotype is in GenBank (Accession XXXXXXXXX).

## Acknowledgments

We thank R. Dalton, A. Freundlich, M. Goldberg, R. Kaczorowski, M. Koski, B. McTeague, G. Meindl, C. Saski, K. Schuller, R. Spigler, L. Stanley, H. Wipf, the University of Pittsburgh GPCL and greenhouse staff, the Oregon State University CGRB, and the University of Idaho IBEST Core Facilities for greenhouse, field, laboratory, or data assistance, and the Ashman and Liston lab members, N. Bassil, and R. Cronn for helpful comments. This work was supported by the National Science Foundation (DEB 1241006 and DEB 1020523 to TLA, and DEB 1241217 and DEB 1020271 to AL), and the UPitt Dietrich School of Arts and Sciences.

## Supporting Information

Figures S1-S3

Tables S1-S5

